## Supplemental Material for "The replicative helicase CMG is required for the divergence of cell fates during asymmetric cell division *in vivo*"

### Supplemental Figures

Memar_FigS1

**A**

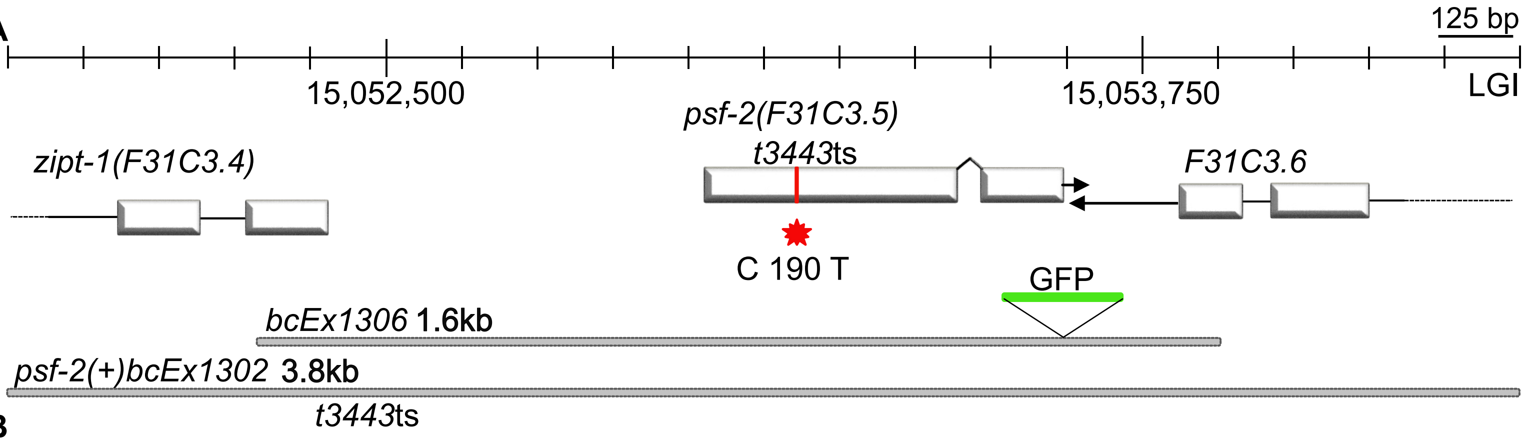


**B**

```
                          P 64 S
Sc      MSLPAHLQQTFSPEEIQFIVENEPIKIFPRITTRQKIRGDDRGTGNHTRWQLITTDDKAL   60
Ce      ----------MNAERCEFIAGNSLIEVIPSISDDRPIHL-------------ISGD---I   34
Dm      ----------MDPSIIEFIGEKCMISIIPNFSN-EPLHL-------------IYGP---V   33
Mm      ----------MDAAEVEFLAEKELVTIIPNFSL-DKIYL-------------IGGD---L   33
Hs      ----------MDAAEVEFLAEKELVTIIPNFSL-DKIYL-------------IGGD---L   33
                  :.    :*:  :  : ::* ::    :            *      :

                                     P 64 S
Sc      NNMVAMRSTEVVLWIALLLKQQSKCSIVAPQWLTTKELDRKIQYEKTHPDRFSELP-WNW   119
Ce      GPFEAGVPCRIPVWTAILMKRKHNCKVVAPQWMDVDELKKI-LTSETESQGLAKLP-DHF   92
Dm      GPFRAGFPVFVPLWMATHLRKQQKCRIVPPEWMDMDILEEI-KEEEKRSKFFTKMPCEHY   92
Mm      GPFNPGLPVDVPLWLAINLKQRQKCRLLPPEWMDVEKLEQM-RDEERKEETFTPVPSPHY   92
Hs      GPFNPGLPVEVPLWLAINLKQRQKCRLLPPEWMDVEKLEKM-RDHERKEETFTPMPSPYY   92
        . :        : :* *  :::: :* :: *:*:  . *..     :  .  . :: :*   :

Sc      LVLARILFNKAKDDFHDPIHELRGKIQDLREIRQIKVLKG----LKYLNESHLQLDNLSL   175
Ce      FEISHMLVRDAREDIFE-VEAVKSLVQDIYDRRDAKLRSSAIEFLRQNQTCHAQLDNVQL   151
Dm      MVVAQLVMSTAPDDVPR-CEELRTVIKDIFDIRESKLRTSIDAFIKGE-GTYAKLDNLTL   150
Mm      MEITKLLLNHASDNIPK-ADTIRTLIKDLWDTRMAKLRVSADSFVRQQ-EAHAKLDNLTL   150
Hs      MELTKLLLNHASDNIPK-ADEIRTLVKDMWDTRIAKLRVSADSFVRQQ-EAHAKLDNLTL   150
        : :::::.   * ::.    . ::  ::*: :  *  *:  .    ::     : :***: *

Sc      LEINELRPFITEIMDKLREIHTASLTAGTENDE------------E---EFNI   213
Ce      IEASSARATLEACRQMGAVV-RNKHESTPL-----------------------   180
Dm      LEIHSVRPILPYSLDHIARYQRTATASQRDTSMLSASMAGSSSGPNSNSLFSQ   203
Mm      MEISSSGAFLTQALNHMYKLRTNLQPSESTQSQ---DF---------------   185
Hs      MEINTSGTFLTQALNHMYKLRTNLQPLESTQSQ---DF---------------   185
        :*      :    :
```

**C**

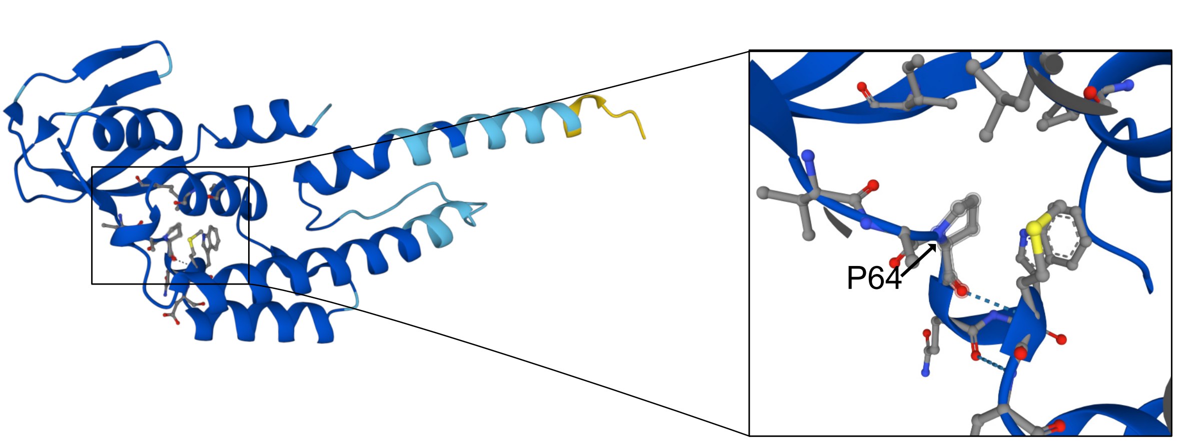

**Supplemental Figure 1. Identification of *t3443*ts mutation in *psf-2* locus and alignment of PSF-2 protein sequence with orthologs in different species. (A)** Top. Schematic of *psf-2* locus and adjacent transcription units on LGI. *t3443*ts is a C-to-T change at position 190bp of *psf-2*'s coding sequence and is indicated in red. Bottom. Schematic of 3.8kb and 1.6kb genomic fragments that were used to generate the rescuing transgenes *bcEx1302* and *bcEx1306*. **(B)** Alignment of PSF-2 protein sequence with orthologs of PSF-2 in different species generated using Clustal Omega[97]. *t3443*ts causes a predicted proline-to-serine change at position 64 of PSF-2's amino acid sequence and is indicated in red. **(C)** Structure of *C. elegans* PSF-2 generated with AlphaFold[98,99]. An enlargement of the region around Proline 64 (P64) is shown on the right.

Memar_FigS2

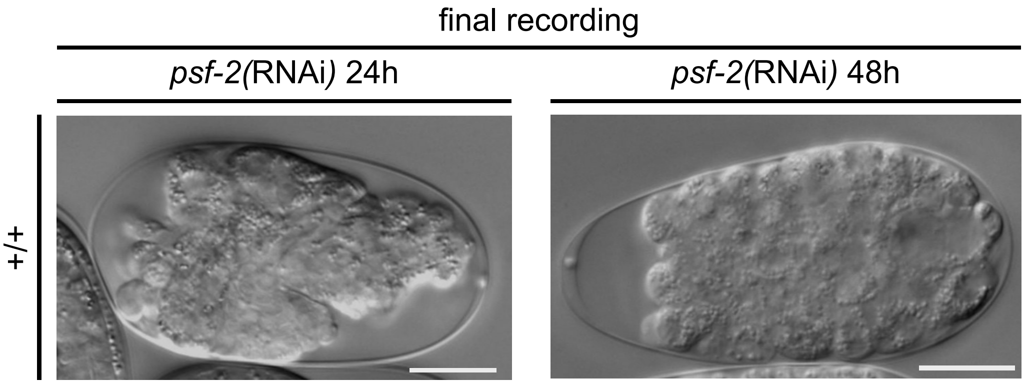

**Supplemental Figure 2. The severity of the *psf-2*(RNAi) phenotype is dependent on the duration of RNAi knock-down.** DIC images of the final recordings for representative embryos. After a 24h RNAi treatment (left), embryos reach and initiate the morphogenesis stage. After a 48h RNAi treatment (right), embryos are unable to reach and initiate the morphogenesis stage. Instead, they arrest after reaching the ~50-cell stage.

Memar_FigS3

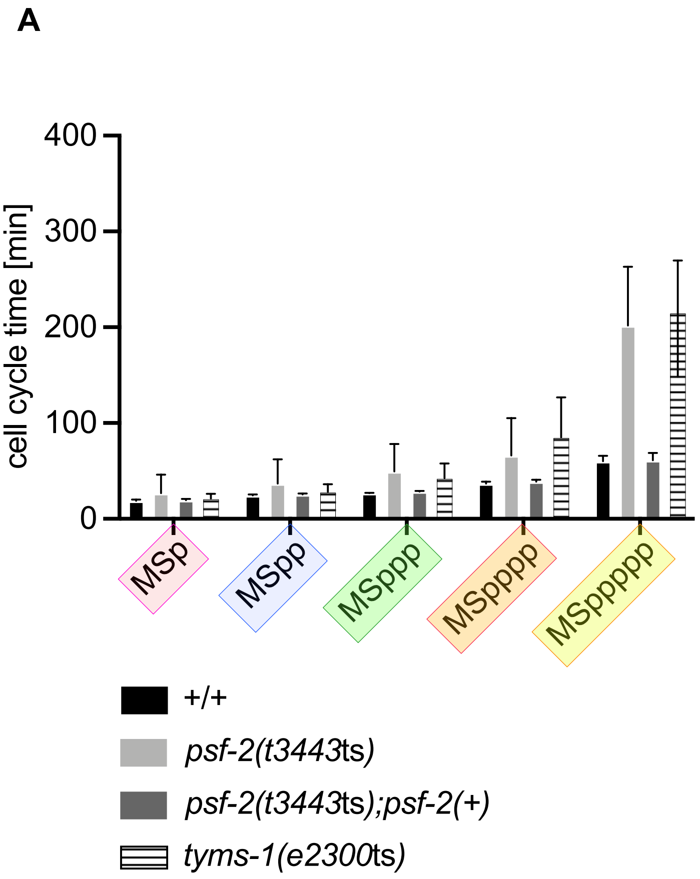
A
400
300
200
100
0
cell cycle time [min]
MSp
MSpp
MSppp
MSpppp
MSppppp
+/+
psf-2(t3443ts)
psf-2(t3443ts);psf-2(+)
tyms-1(e2300ts)

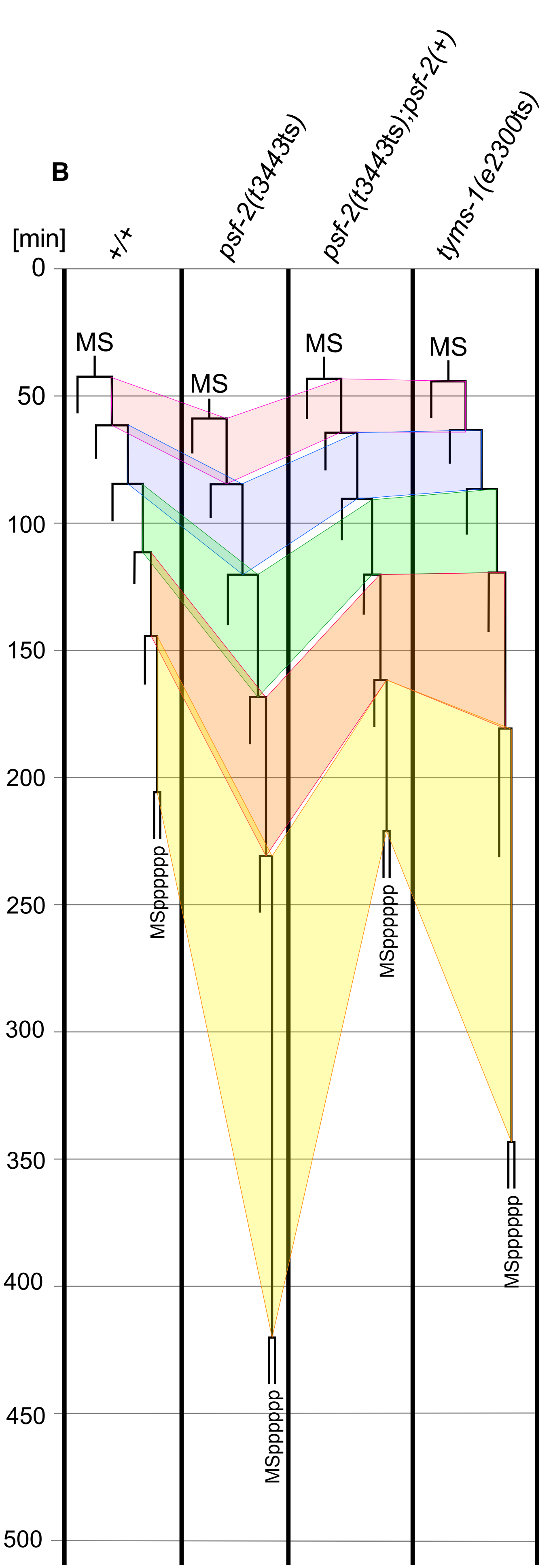
B
[min]
+/+
psf-2(t3443ts)
psf-2(t3443ts);psf-2(+)
tyms-1(e2300ts)
0
50
100
150
200
250
300
350
400
450
500
MS
MS
MS
MS
MSpppppp
MSpppppp
MSpppppp
MSpppppp

**Supplemental Figure 3. The increase in cell cycle length in *psf-2(t3443*ts) is not lineage-dependent. (A)** Cell cycle length [min] of the MSppppp cell and its ancestors in wild-type (+/+) (n=6), *psf-2*(*t3443*ts) (n=6), *psf-2*(*t3443*ts); *psf-2*(+) (transgene *bcEx1302*) (n=5) and *tyms-1*(*e2300*ts) (n=5) animals at 25°C. Mean ±SD are indicated. (B) MSpppppp lineage of representative animals of the genotypes indicated. The wild-type and *psf-2*(*t3443*ts); *psf-2*(+) embryos analyzed completed embryogenesis and hatched. Analyses were performed at 25 °C and *tyms-1(e2300*ts*)* embryos were shifted to 25 °C at the 1-4-cell stage.

Memar_FigS4

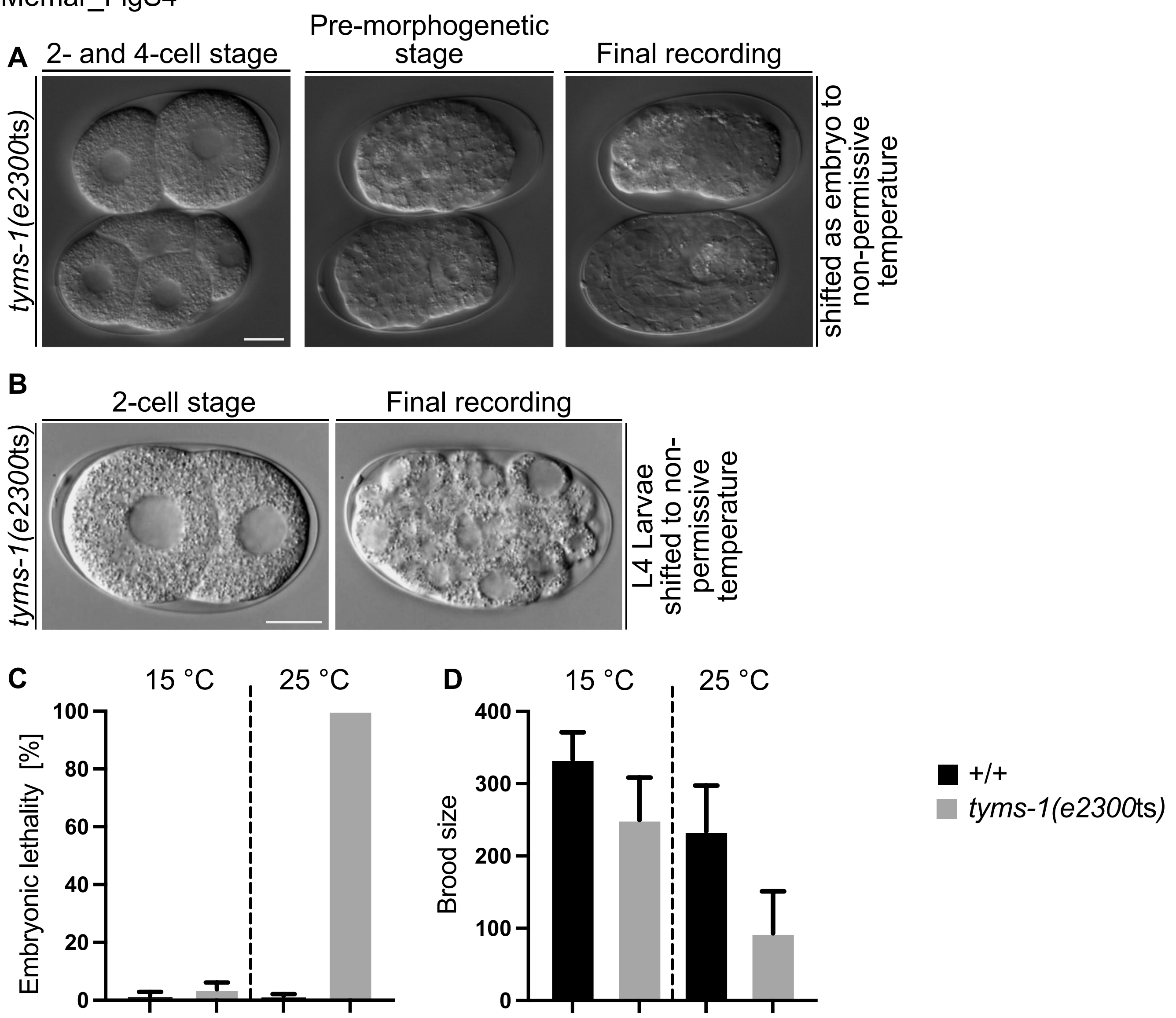


**E**

```
Sc --------------------------MTMDGKNKEEEQYLDLCKRIIDEGEFRPDRTGTGTLSLFAPPQLRFSLRDDTFPLLTTKK 60
Ce ----MN---KENI-IADAPSDVVKTVQQQVHLNQDEYKYLKQVEQILREGTRRDDRTGTGTISIFG-MQSKYCLRNGTIPLLTTKR 77
Dm MVLTPTKDGPDQESMPLPADNGESPSKQQAPVNRDEMHYLDLLRHIIANGEQRMDRTEVGTLSVFG-SQMRFDMRN-SFPLLTTKR 84
Mm ----MLVVGSELQSDA------QQL--SAEAPRHGELQYLRQVEHILRCGFKKEDRTGTGTLSVFG-MQARYSLRD-EFPLLTTKR 72
Hs ----MPVAGSELPRRPLPPAAQERD--AEPRPPHGELQYLGQIQHILRCGVRKDDRTGTGTLSVFG-MQARYSLRD-EFPLLTTKR 78
                                         : * :**     .:*:  *  : *** .**:*:*.  * :: :*:  :******:
```

W 80 C

```
Sc VFTRGIILELLWFLAGDTDANLLSEQGVKIWDGNGSREYLDKMGFKDRKVGDLGPVYGFQWRHFGAKYKTCDDDYTGQGIDQLKQV 146
Ce VYWKGVLEELLWFISGSTDGKLLMEKNVKIWEKNGDRAFLDNLGFTSREEGDLGPVYGFQWRHFGAKYVDCHTDYSGQGVDQLAEV 163
Dm VFFRAVAEELLWFVAGKTDAKLLQAKNVHIWDGNSSREFLDKMGFTGRAVGDLGPVYGFQWRHFGAQYGTCDDDYSGKGIDQLRQV 170
Mm VFWKGVLEELLWFIKGSTNAKELSSKGVRIWDANGSRDFLDSLGFSARQEGDLGPVYGFQWRHFGAEYKDMDSDYSGQGVDQLQKV 158
Hs VFWKGVLEELLWFIKGSTNAKELSSKGVKIWDANGSRDFLDSLGFSTREEGDLGPVYGFQWRHFGAEYRDMESDYSGQGVDQLQRV 154
    *: :.:  *****: *.*:.: *  :.*:**: *..* :**.:**. *  ****************:*     .**:*:*:*** .*
Sc IHKLKTNPYDRRIIMSAWNPADFDKMALPPCHIFSQFYVSFPKEGEGSGKPRLSCLLYQRSCDMGLGVPFNIASYALLTRMIAKVV 233
Ce IRQIKEQPDSRRIIMSAWNPSDLGQMVLPPCHTMCQFYVD---------NGELSCQLYQRSGDMGLGVPFNLASYGLLTHMIAKVC 241
Dm IDTIRNNPSDRRIIMSAWNPLDIPKMALPPCHCLAQFYVSEK-------RGELSCQLYQRSADMGLGVPFNIASYALLTHMIAHVT 250
Mm IDTIKTNPDDRRIIMCAWNPKDLPLMALPPCHALCQFYVV---------NGELSCQLYQRSGDMGLGVPFNIASYALLTYMIAHIT 236
Hs IDTIKTNPDDRRIIMCAWNPRDLPLMALPPCHALCQFYVV---------NSELSCQLYQRSGDMGLGVPFNIASYALLTYMIAHIT 232
    *  :: :*  .*****.****   *:  *.*****
Sc DMEPGEFIHTLGDAHVYKDHIDALKEQITRNPRPFPKLKIKRDVKDIDDFKLTDFEIEDYNPHPRIQMKMSV 304
Ce GLKPGTLVHTLGDAHVYSNHVDALKIQLDREPYAFPKIRFTRDVASIDDFTSDMIALDDYKCHPKIPMDMAV 312
Dm GLKPGDFVHTMGDTHVYLNHVEPLKEQLERTPRPFPKLIIKRQVQDIEDFRFEDFQIVDYNPHPKIQMDMA- 320
Mm GLQPGDFVHTLGDAHIYLNHIEPLKIQLQREPRPFPKLKILRKVETIDDFKVEDFQIEGYNPHPTIKMEMAV 307
Hs GLKPGDFIHTLGDAHIYLNHIEPLKIQLQREPRPFPKLRILRKVEKIDDFKAEDFQIEGYNPHPTIKMEMAV 313
    .::** ::**:**:*:* :*:: ** *: * *      ***: : *.*  *:**    : : .*: ** * *.*:
```

**Supplemental Figure 4. Identification of *e2300*ts mutation in *tyms-1* locus and alignment of TYMS-1 protein sequence with orthologs in different species.** (**A**) DIC images of representative *tyms-1*(*e2300*ts) embryos at the 2- and 4-cell stage, the pre-morphogenetic stage and at the final recording (terminal phenotype). Embryos were shifted to 25°C at the 2- to 4-cell stage. (**B**) DIC images of representative *tyms-1*(*e2300*ts) embryos at the 2-cell stage and at the final recording (terminal phenotype). L4 larvae were shifted to 25°C, 2-cell stage embryos were extracted after 16h and mounted for long-term imaging. For both (A) and (B), images were taken from long-term recordings performed at 25°C. Scale bars represent 10 μM. **(C)** Embryonic lethality [%] and **(D)** Brood size at permissive (15°C) and non-permissive temperature (25°C) in wild-type (+/+) and *tyms-1*(*e2300*ts). Embryonic lethality and brood size of four (n=4) or five (n=5) adults were analyzed in the case of wild-type (15°C or 25°C, respectively). In the case of *tyms-1*(*e2300*ts), embryonic lethality and brood size of five adults (n=5) were analyzed. Mean ±SD are indicated. **(E)** Alignment of the TYMS-1 protein sequence with orthologs of thymidylate synthetase in different species generated using Clustal Omega[97].

Memar_FigS5

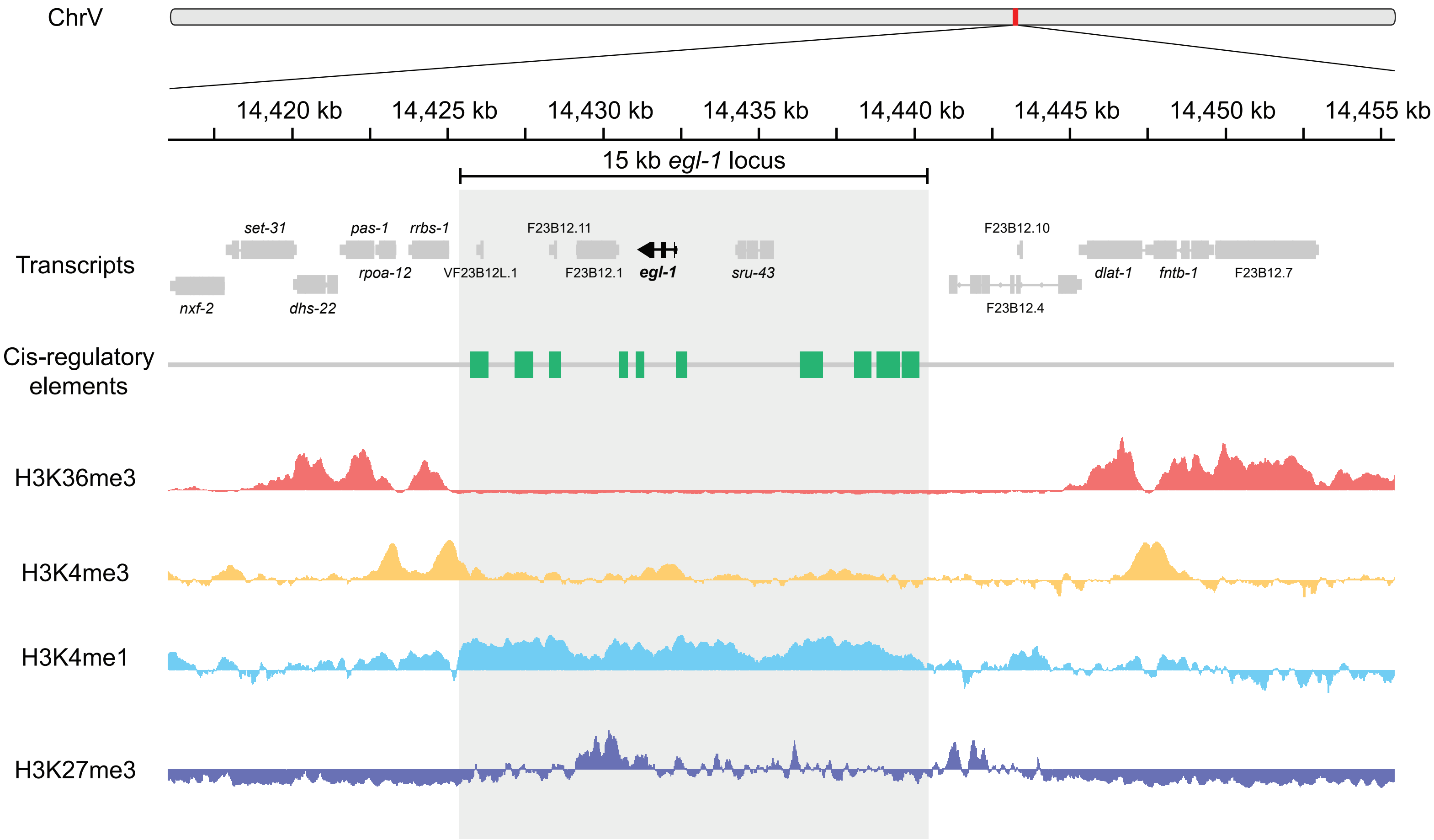

ChrV
14,420 kb
14,425 kb
14,430 kb
14,435 kb
14,440 kb
14,445 kb
14,450 kb
14,455 kb
15 kb egl-1 locus
Transcripts
set-31
pas-1
rrbs-1
F23B12.11
F23B12.10
nxf-2
dhs-22
rpoa-12
VF23B12L.1
F23B12.1
egl-1
sru-43
dlat-1
fntb-1
F23B12.7
F23B12.4
Cis-regulatory elements
H3K36me3
H3K4me3
H3K4me1
H3K27me3

**Supplemental Figure 5. Genome tracks of ChrV:14,415,000-14,455,000 (40 kb)**

Within the 40 kb view, the 15 kb *egl-1* locus containing the transcript and cis-regulatory elements is highlighted in grey. All transcripts (www.wormbase.org)[89,90] within the 40 kb locus are indicated in grey, except the *egl-1* transcript, which is indicated in black. The *cis*-regulatory elements of the *egl-1* locus are indicated in green[18]. ChIP-seq peaks obtained from bulk *C. elegans* embryos for various histone H3 modifications were mined from publicly available datasets[70]. H3K36me3 tracks are indicated in red, H3K4me3 tracks are indicated in yellow, H3K4me1 tracks are indicated in light blue and H3K27me3 tracks are indicated in dark blue. This figure was adapted from the Integrative Genomics Viewer Application https://igv.org/app/[100].
